# Automated classification of cellular expression in multiplexed imaging data with Nimbus

**DOI:** 10.1101/2024.06.02.597062

**Authors:** J. Lorenz Rumberger, Noah F. Greenwald, Jolene S. Ranek, Potchara Boonrat, Cameron Walker, Jannik Franzen, Sricharan Reddy Varra, Alex Kong, Cameron Sowers, Candace C. Liu, Inna Averbukh, Hadeesha Piyadasa, Rami Vanguri, Iris Nederlof, Xuefei Julie Wang, David Van Valen, Marleen Kok, Travis J. Hollmann, Dagmar Kainmueller, Michael Angelo

## Abstract

Multiplexed imaging offers a powerful approach to characterize the spatial topography of tissues in both health and disease. To analyze such data, the specific combination of markers that are present in each cell must be enumerated to enable accurate phenotyping, a process that often relies on unsupervised clustering. We constructed the Pan-Multiplex (Pan-M) dataset containing 197 million distinct annotations of marker expression across 15 different cell types. We used Pan-M to create Nimbus, a deep learning model to predict marker positivity from multiplexed image data. Nimbus is a pre-trained model that uses the underlying images to classify marker expression across distinct cell types, from different tissues, acquired using different microscope platforms, without requiring any retraining. We demonstrate that Nimbus predictions capture the underlying staining patterns of the full diversity of markers present in Pan-M. We then show how Nimbus predictions can be integrated with downstream clustering algorithms to robustly identify cell subtypes in image data. We have open-sourced Nimbus and Pan-M to enable community use at https://github.com/angelolab/Nimbus-Inference.

## Introduction

Recent developments in instrumentation have made highly multiplexed protein imaging more routine, with multiple mass spectrometry and optical microscopy platforms capable of measuring 10s to 100s of proteins on large, intact tissue sections^1–4^. This increase in throughput and multiplexing have added a spatial domain to the single-cell revolution, unlocking the ability to catalogue the full complement of cells present in a sample, understand their spatial organization, and infer their interactions. These techniques have proven invaluable in understanding how structure and function are interrelated in tissue homeostasis, the tumor microenvironment, and during infection^5–7^. This deluge of data has necessitated the development of algorithms across the full spectrum of the analysis pipeline to translate the raw imaging measurements into biological insights.

Cell type assignment is a crucial step in the analysis and interpretation of single-cell spatial data. This core step is shared across nearly all single cell technologies—including flow cytometry, mass cytometry, single cell RNA-seq, and single cell ATAC-seq. Although the approach, data modality, and biomolecules of interest can vary significantly, the end goal is the same: to assign cells to a cell type based on the combinatorial expression of the detected biomolecules. In line with the importance of this task, substantial effort has been devoted to developing more robust and automated methods for cell type assignment. These include approaches based on decision trees, hierarchical clustering, self-organizing maps, unsupervised graph-based clustering, and mapping to reference atlases^8–16^.

Although single cell information can be extracted from spatial data, there is a key difference from other single cell techniques—there is no dissociation step that physically separates adjacent cells from one another. Thus, images of intact tissue are not inherently single cell when generated. Instead, cells in the image must be identified through a process known as cell segmentation, where the border of individual cells is detected, labeled, and disambiguated from overlapping and adjacent cells. There are now deep learning models that can generate high-quality cell segmentations with human-level accuracy for most tissue types ^17–19^. Once cell segmentation labels have been generated, the co-occurrence of protein or RNA expression within each cell can be quantified. This is typically calculated by averaging the intensity of a given marker across all pixels within the cell, otherwise known as integrated expression.

Despite these advances, cell segmentation using software or by a human is rarely perfect, and some degree of error is inevitable. Even with perfect segmentation, multiple confounders inherent to imaging data make accurate cell type assignments a challenge in two-dimensional imaging data. For example, shared boundaries between bordering cells can cause signal to spill over into adjacent cells, especially for markers localized to the cell membrane. Furthermore, tissues can have background staining that does not represent biological signal, such as autofluorescence or non-specific staining. Additionally, marker intensities can vary over several orders of magnitude such that universal cut points for assigning marker positivity cannot be used. As a result, simply averaging the expression across all the pixels within a cell, via integrated expression, is often an unreliable proxy for determining cell positivity. Most of these confounding factors are readily apparent to trained experts (i.e., pathologists) and can be taken into account during manual scoring. The subcellular pattern, intensity, and contrast of marker expression with respect to its nearby surroundings provide a spatial context that is invaluable for determining whether a cell is positive for a given proteomic marker. However, manually scoring cells in highly multiplexed imaging data is not scalable. As a result, nearly all existing algorithms use integrated expression for cell type assignment^10,13–16,20,21^. This simplification has major benefits in generalization, computational efficiency, and interoperability for algorithm developers, and has been the natural choice in the absence of viable alternatives. Unfortunately, this choice results in the loss of critical spatial information that could greatly enhance the accuracy of cell type assignment.

Convolutional Neural Networks (CNNs) are a form of deep learning that have achieved human-level accuracy across a wide range of challenging domains in biological imaging^17,22^, including super-resolution imaging^23^, spot detection^24^, image denoising^18^, cell segmentation^17,19^, and disease classification^25,26^. CNNs are appealing because they are trained to make predictions directly using the original image as an input. Model training typically requires a labeled dataset with many examples of the task the algorithm will perform in order to learn how to make valid predictions without overfitting. This presents two key challenges for training a deep learning algorithm for cell classification. First, manual cell type annotation is laborious and requires significant expertise. Second, models trained for direct cell type prediction^20,21,27^ will only be valid for the specific set of markers included in their training data, limiting generalization to other datasets where markers might differ.

Here, we set out to create a single deep learning model for human-like, visual classification of marker positivity that would generalize across tissue types, image platforms, and markers. To overcome the challenges outlined above that are inherent to training a deep learning model for direct cell type prediction, we instead split the task into two separate steps. We first leveraged previously published and unpublished multiplexed proteomic imaging datasets to create the Pan-Multiplex dataset (Pan-M, **Fig. 1c**), which contains more than 197 million annotations across 56 proteins and 10 cell lineages (**Fig. 1b**). We used Pan-M to train Nimbus, a deep learning model that predicts marker positivity independently for each channel (**Fig. 1a**), overcoming the limitations of integrated expression. We then used the predictions generated by Nimbus, instead of integrated expression, as inputs to conventional clustering algorithms (**Fig. 1c**), showing how this workflow achieves accurate cellular phenotyping without laborious manual scoring or expert-level correction. Importantly, Nimbus can be run on any multiplexed antibody dataset (without finetuning or retraining) to generate accurate single-cell predictions that are robust to the confounders that affect integrated expression. This approach addresses the root cause that makes cell clustering more difficult with spatial data relative to dissociated single-cell assays. We have open-sourced both the Pan-M dataset and Nimbus to serve as useful tools for the community.

**Fig. 1.**
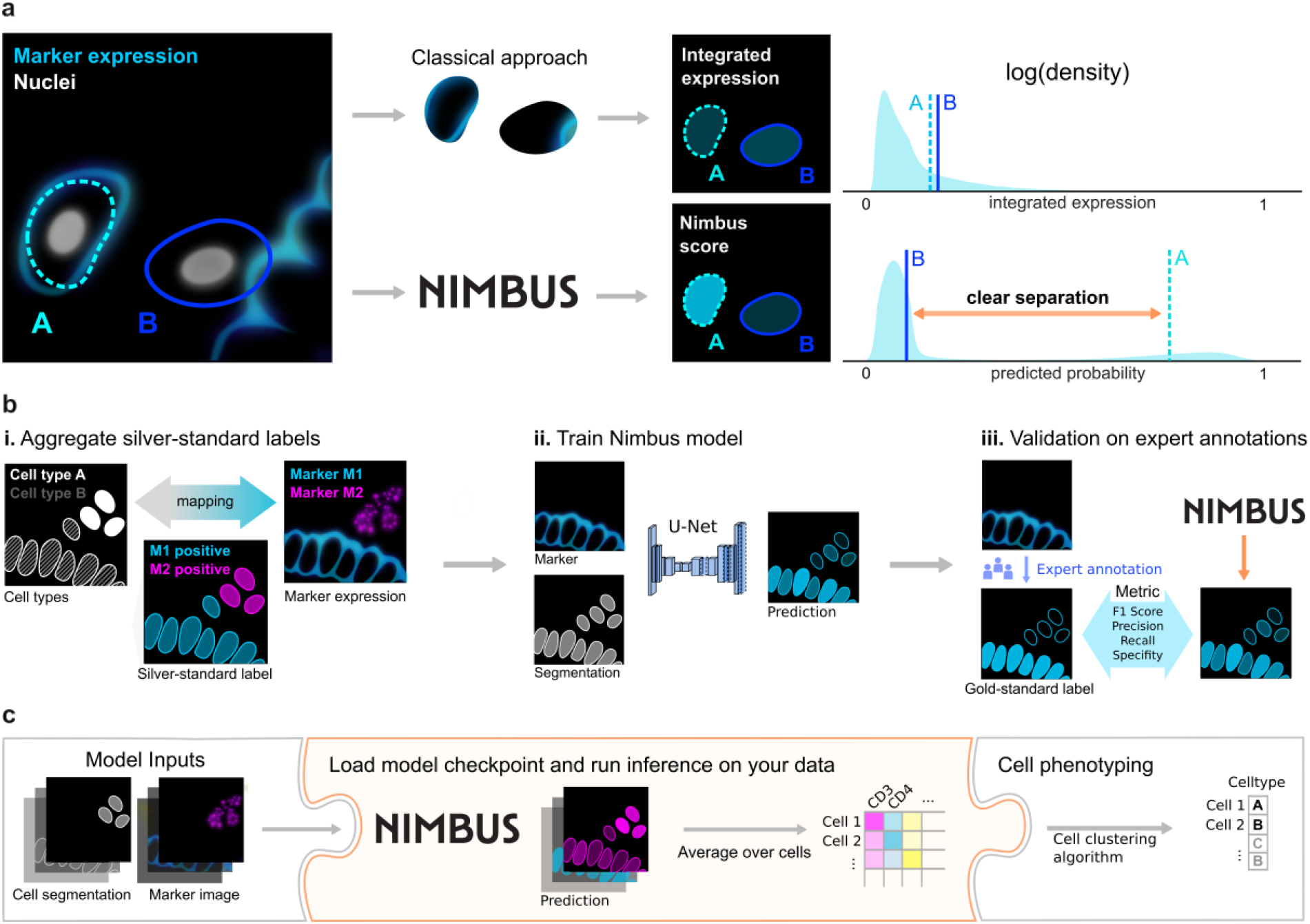
NIMBUS improves marker prediction for phenotyping in multiplex images. **a**, Nimbus enables better separation between positive and negative cells by incorporating subcellular expression patterns, compared to integrated expression which just uses the average. **b**, The Nimbus model is a noise-robust U-Net trained on a diverse set of publicly available multiplexed imaging datasets (silver standard labels) with subsequent expert validation (gold standard labels). **c**, Drop-in integration of Nimbus in various cell phenotyping pipelines.

## Main

### Constructing the Pan-M dataset

Deep learning models require large amounts of labeled training data. This volume of data is needed in order to prevent overfitting, and underpins the success of recent efforts to predict protein structures, identify transcription factor binding sites, and segment cells^17,28,29^. Our goal was to construct a dataset which would facilitate the training of accurate deep learning models to predict marker positivity on a single-cell basis. Given the pivotal role that training data plays in enabling accurate models, it is crucial to construct a training dataset that captures the breadth and diversity of data that the final trained model will be run on to ensure that it makes accurate predictions.

To create a sufficiently large and diverse dataset, we built a pipeline to extract training data from published^5,30^ and unpublished multiplexed imaging datasets where the cells had been clustered using conventional approaches. For each image, we collated 1) the segmentation mask, which denotes the precise location and shape of every cell, 2) the table of cell assignments, which labels every cell with its cell type, and 3) the individual channels of imaging data. Based on the cell type assignments, we then generated an assignment matrix, which mapped cell types to channel positivity (**Fig. 2a and Extended Data Fig. 1 a-c**). For example, CD8T cells would be marked as positive for CD3, CD8, and CD45, whereas CD4T cells would be marked as positive for CD3, CD4, and CD45. This was done for each cell type in the dataset, across all of the channels used for clustering. This assignment matrix was then used to produce the silver standard labels (**Fig. 2a**). We refer to them as silver standard labels because they depend on the accuracy of the initial clustering, rather than any manual proofreading.

**Fig. 2.**
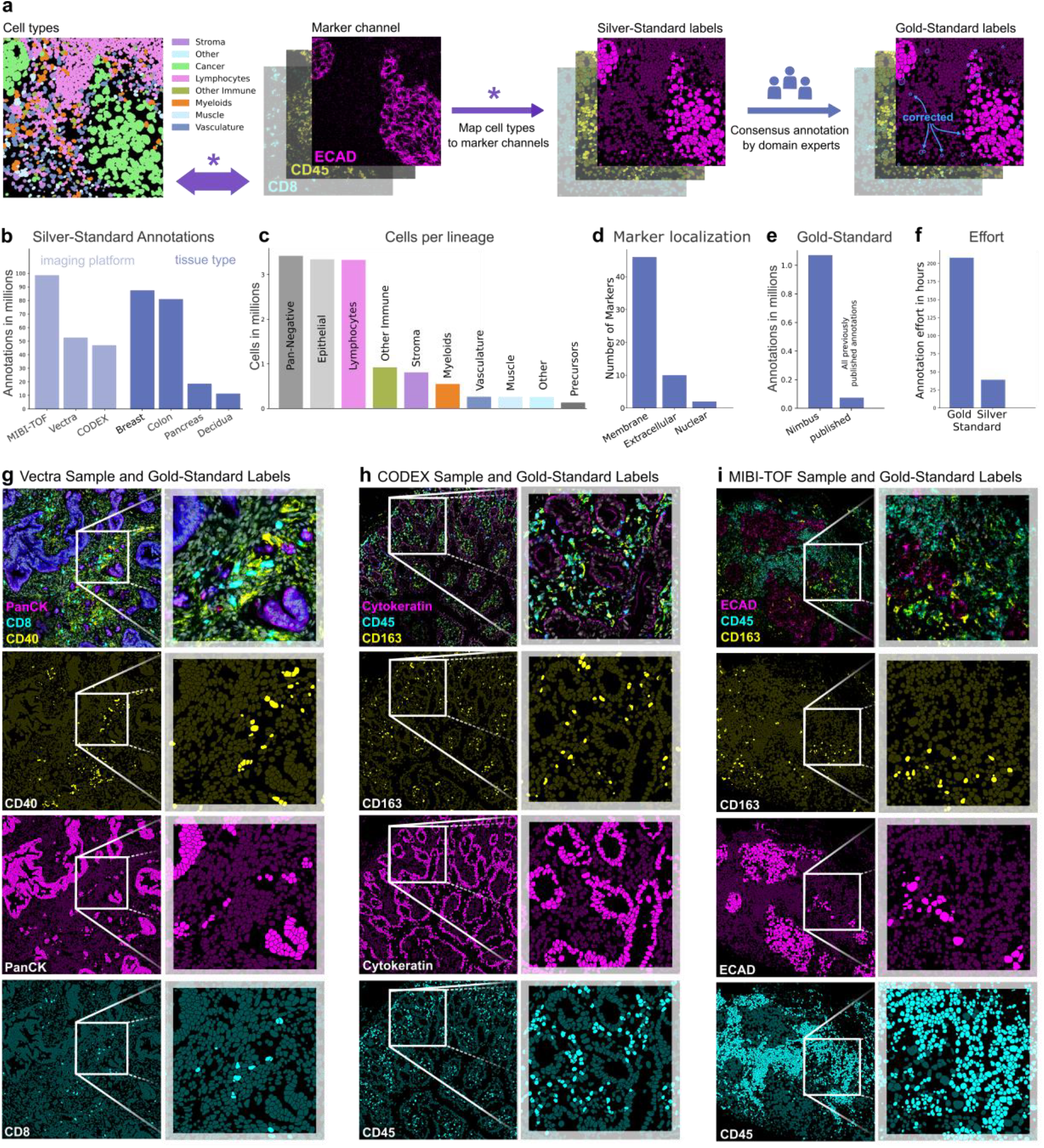
Data Annotation. **a**, Schematic of the pipeline to generate the Pan-M dataset. We mapped single channel positivity from previously clustered data to produce the silver standard labels. We then manually curated a subset of these images to generate gold standard labels. **b**, The number of annotations across image platforms and tissue types. **c**, The number of cells of each cell type. **d**, The subcellular localization of the included markers. **e**, The number of gold standard annotations in Pan-M compared to previously published. **f**, The number of hours required to generate the gold standard and silver standard labels. **g-i**, Image samples with corresponding gold standard labels for the imaging platforms Vectra, CODEX and MIBI-TOF, respectively.

We used our silver standard label pipeline to generate Pan-M, which contains 197 million annotations across 15 million cells. The Pan-M dataset includes images from three different image platforms and four different tissue types (**Fig. 2b**). In addition, it contains a diversity of cell types (**Fig. 2c**) spanning epithelial, immune, and stromal populations, as well as a substantial fraction of cells that are negative for all of the included imaging markers (pan-negative). The dataset contains 56 distinct protein markers with a range of staining patterns (**Fig. 2d)**.

Following generation of the silver standard labels for the Pan-M dataset, we selected a subset of images and generated gold standard labels (**Fig. 2a**). These labels were created via manual correction of the silver standard labels by directly examining the multiplexed images to confirm cell positivity for each channel in each cell. In total, we generated over 1 million gold standard annotations, which is significantly more than all of the previously published manually curated annotations for cell type assignment combined (**Fig. 2e**). Although the gold standard annotations are higher quality, they are also much more labor intensive to generate. Each cell in an image must be manually inspected, which scales linearly with the number of proofread cells. In contrast, the silver standard labels can be generated far more efficiently; once the assignment matrix for a given dataset is proofread, it can be applied across all of the cells. As a result, generating gold standard labels takes approximately 900 times longer for the same number of annotations (**Fig. 2f**), which is why we manually annotated only a small subset of the cells in the Pan-M dataset. In **Figs. 2g-i**, we highlight representative images of the gold standard annotation across the three microscopy platforms in Pan-M.

### Nimbus assessment

After constructing the Pan-M dataset, we used it to train Nimbus—a deep learning model to directly predict cell marker positivity. Nimbus is built off of the U-Net architecture^31,32^, which was designed for biomedical image data to capture both high-level features and local details. The inputs to Nimbus are a segmentation mask and a single channel of image data. The output is a score for each cell in the image, ranging from 0 to 1, corresponding to whether that cell is positive for the supplied marker (**Fig. 1a**). We intentionally designed Nimbus to have a simple workflow, with only a single channel of image data required to make predictions. As a result, Nimbus can be run on any multiplexed dataset with any combination of markers, since each marker is treated independently. This is in contrast to previous deep learning algorithms for cell classification^20,21,27^ which require the model to be retrained, since they have learned the specific mapping between markers and cell types present in each dataset.

Nimbus was trained on the silver standard labels in Pan-M (see Methods), which were derived from cluster assignments generated by the original study authors using previous approaches for cell clustering. This is a laborious process which often involves comparisons of different clustering algorithms, multiple rounds of optimization and parameter fine-tuning, along with substantial manual intervention and adjustment in order to get sufficiently accurate cell labels. As a result, generating accurate clustering can take weeks for large datasets. In contrast to this bespoke approach, we trained a single Nimbus model using the same settings across all the underlying data at once. Across the five distinct datasets which make up Pan-M, Nimbus generates predictions which correspond visually to the underlying data (**Fig. 3**). Nimbus accurately identifies marker positivity across different datasets, tissue types, and channels, producing output that corresponds with the underlying shape and structure of the data. For example, Nimbus correctly identifies concentric layers of smooth muscle (SMA+) and endothelium (CD31+) in decidual spiral arteries (**Fig. 3a**), as well as scattered CD45+ immune cells in the surrounding tissue. In the colon (**Fig. 3c**), Nimbus is able to demarcate the MUC1+ epithelial cells from the surrounding VIM+ stromal cells, as well as CD4+ immune cell aggregates.

**Fig. 3.**
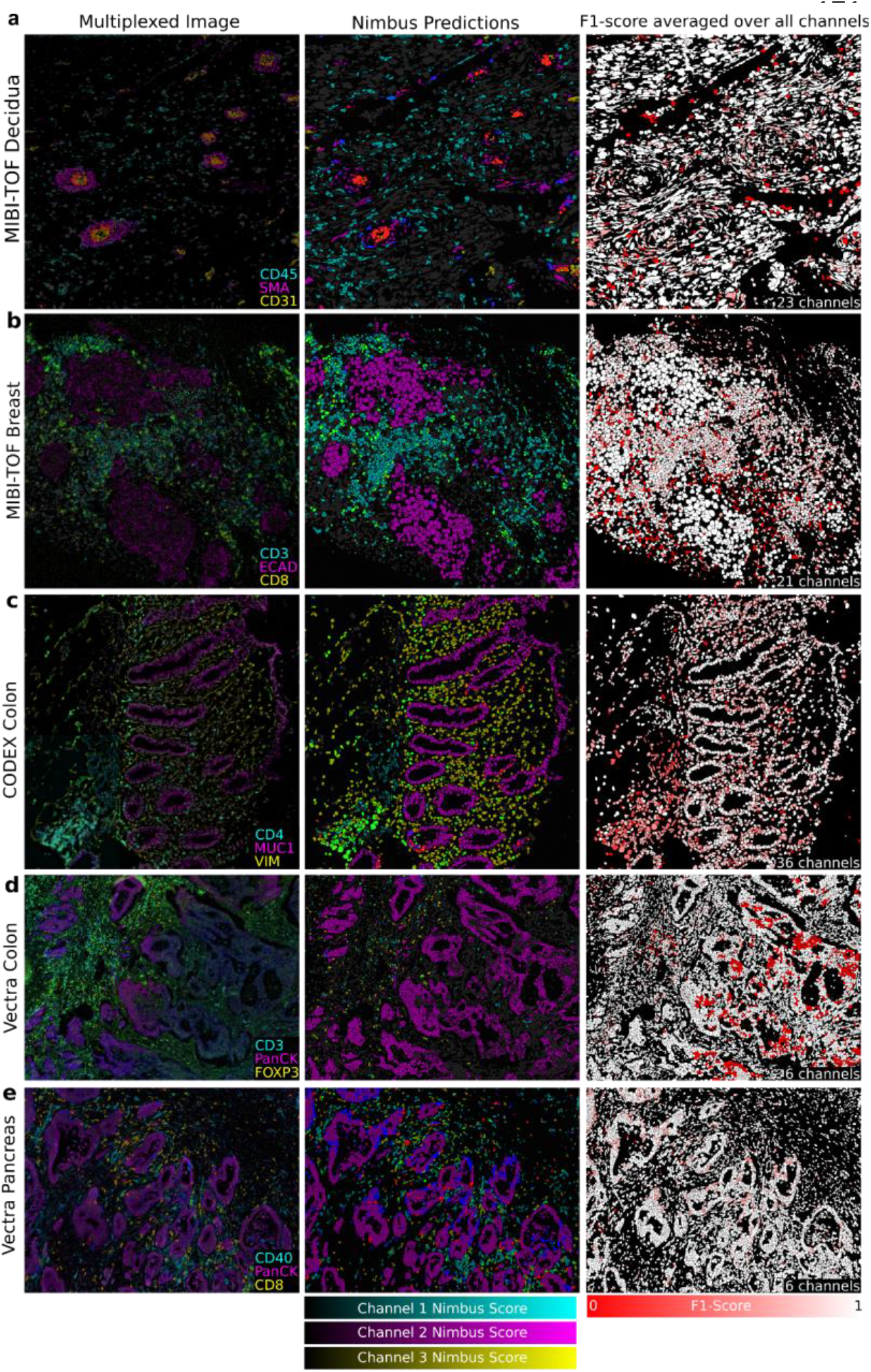
Qualitative Evaluation. **a-e**, Representative images across the five datasets in Pan-M showcasing Nimbus predictions. The left column contains three distinct channels from each dataset. The middle column shows Nimbus predictions for the same three channels, pseudo-colored to align with the color of the imaging channel. The right column shows F1208 scores averaged over all markers.

Moving beyond a qualitative assessment, we next systematically evaluated the accuracy of the Nimbus predictions. We used the gold standard annotations from the held-out test set as our ground truth, comparing the accuracy of the Nimbus scores as well as the original clustering using a number of distinct metrics (see Methods). Across each of these metrics, we see that the Nimbus predictions on the gold standard labels are as accurate as the silver standard labels (**Fig. 4a**). This is true across the different tissue types present in Pan-M (**Fig. 4b**), and across nearly every cell type as well (**Fig. 4c**). Thus, Nimbus represents a single, pre-trained deep learning model for marker classification with accuracy that matches each of the individual clustering solutions employed by the different study authors in the datasets that went into Pan-M.

**Fig. 4.**
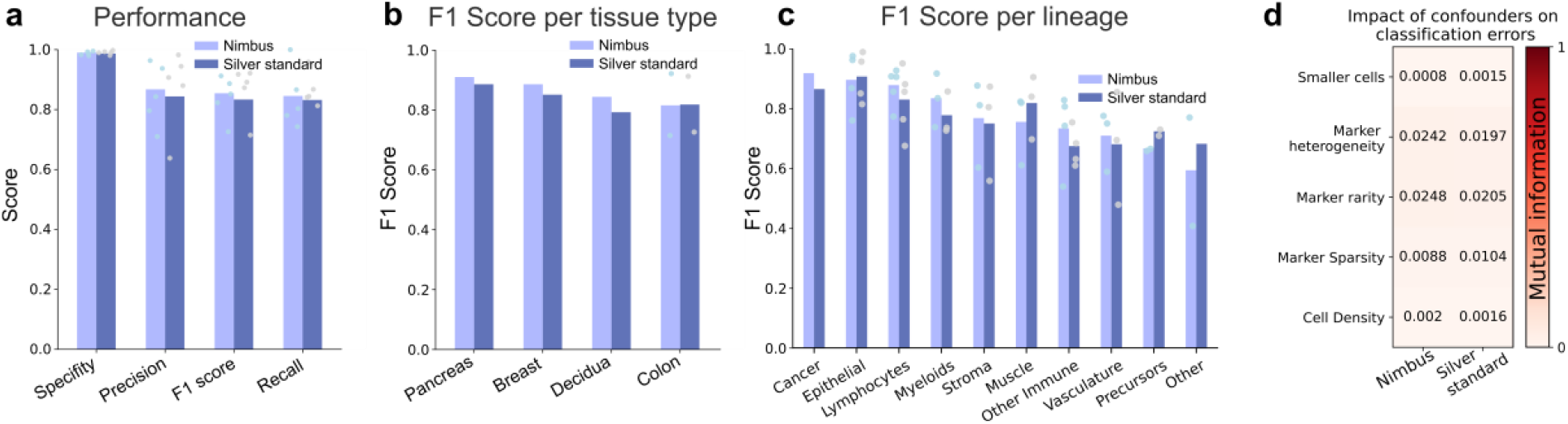
Quantitative Evaluation. **a**, Performance metrics of the Nimbus model and the silver standard labels are compared to gold standard annotations **b-c**, Performance metrics split by tissue type and cell type. **d**, The effect of confounders on the model performance in terms of mutual information.

We next sought to identify factors that impacted model performance. We calculated metrics to define cell density, cell size, marker heterogeneity, and marker sparsity across the test set (see Methods). We then assessed how these features impacted the accuracy of the model. Overall, we observed little impact on performance for the confounders we measured (**Fig. 4d**), suggesting that Nimbus will be able to generalize beyond the specific data it was trained on. Looking at the model architecture itself, we tested the impact of changing the backbone, changing the resolution of the input data, and changing the training schema, none of which significantly affected performance (**Extended Data Fig 1d-g, Supplementary Table 1**).

### Nimbus scores enable accurate cellular phenotyping

Having shown that Nimbus generalizes across datasets, tissues, and cell types to predict marker expression, we next sought to show the advantages of using Nimbus-derived estimates of marker positivity. As discussed in the Introduction, nearly all algorithms developed for clustering image data take the average value of each marker in each cell, which we refer to as integrated expression, as their input. Although easy to compute and convenient to work with, using integrated expression as an input necessarily discards spatial information present for a given marker.

To demonstrate the improvement that Nimbus scores represent over integrated expression, we analyzed the distributions of both metrics in the gold standard test set. Across all channels in all images, Nimbus scores showed clear separation between the gold standard positive and negative populations (**Fig. 5a, left**), indicating that a higher Nimbus score was a reliable proxy for true cell positivity. In contrast, integrated expression did not exhibit the same pattern (**Fig. 5a, right**). In particular, due to the challenges of capturing complex spatial information with a simple average, there was substantial overlap between the gold standard positive and negative populations. Studying a specific channel, we saw the same pattern with Cytokeratin; the Nimbus scores were well-separated between the gold standard positive and negative cells with two almost completely non-overlapping distributions of predicted positivity for Cytokeratin (**Figure 5b, left**). In contrast, when looking at integrated expression, the distributions overlapped substantially (**Figure 5b, right**).

**Fig. 5.**
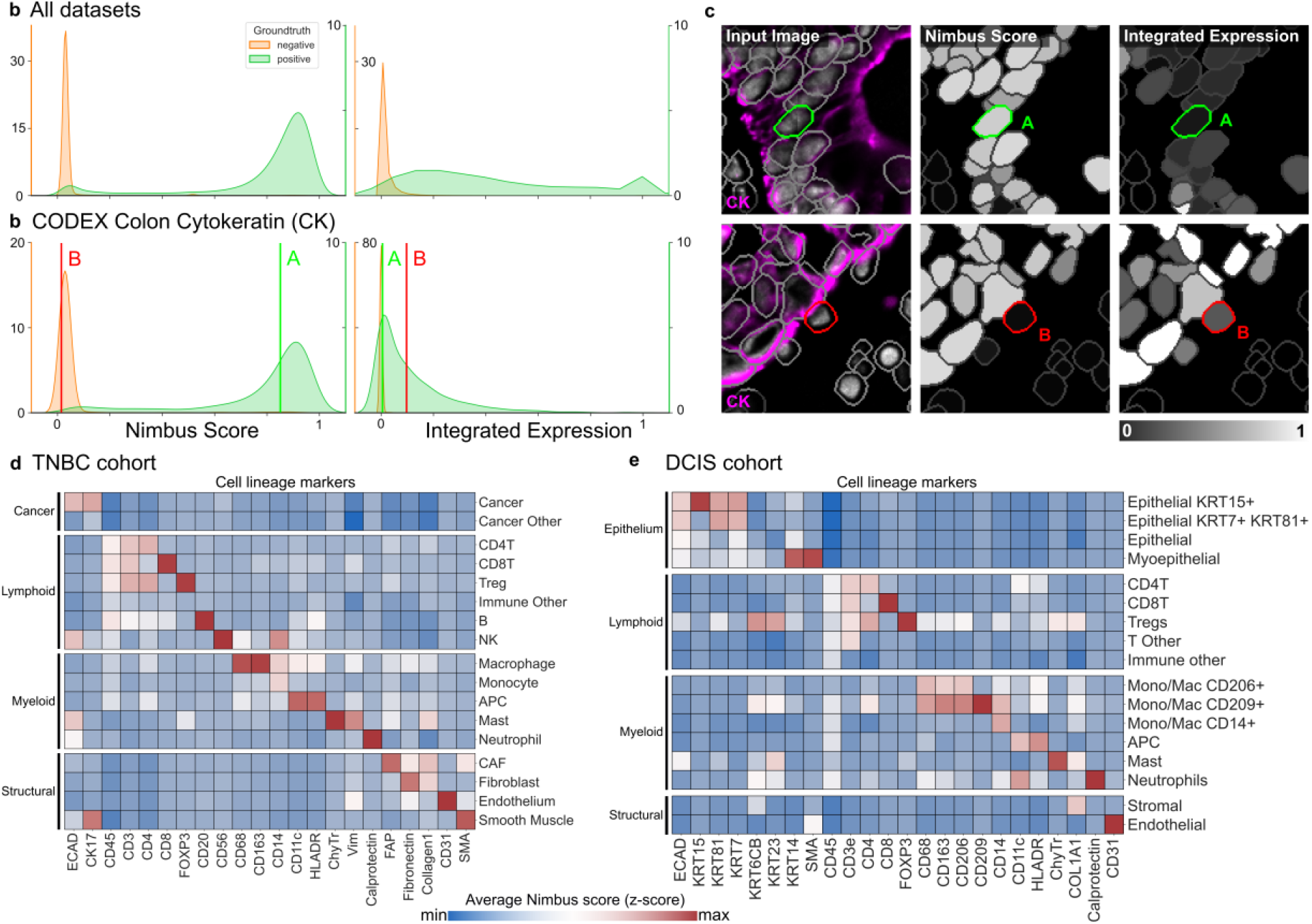
Bi-modality of Nimbus confidence scores. **a**, Kernel density estimate plots of the distribution of Nimbus confidence scores on the left and integrated expression on the right for all gold standard annotated cells. **b**, Kernel density estimate plot of the Nimbus scores and integrated expression for the Cytokeratin channel in a CODEX colon cancer dataset shown in **c. d-e** Cellular phenotypes assigned based on Nimbus scores for a TNBC and a DCIS cohort, which were not part of the training data.

To highlight why Nimbus outperformed integrated expression, we identified specific cells that exemplify the source of this discrepancy. When marker staining is dim, and located at the periphery of a cell, the value of integrated expression will be quite low. However, visually inspecting the image shows that even though the signal contained with the segmented cell is low, the cell is indeed positive for that marker. This is what is highlighted in the top row of **Fig. 5c**, where Nimbus accurately identified positive expression in a cell with low integrated expression. Alternatively, when very bright signal from one cell spills over into an adjacent cell, the integrated expression resulting from that spillover can be quite high. However, inspecting the image demonstrates that this cell is not actually positive for the marker, it is simply in close physical proximity to the cell with the real signal. This is highlighted in the bottom row of **Fig. 5c**, where Nimbus accurately identified negative expression of Cytokeratin in the highlighted cell despite a high value for integrated expression.

Given that Nimbus scores better delineate positive and negative cells from one another compared to integrated expression, we hypothesized that using these scores instead of integrated expression would make unsupervised clustering significantly faster and require less manual adjustment and finetuning. To demonstrate this, we first generated Nimbus scores for all markers and cells in a Multiplexed Ion Beam Imaging (MIBI) breast cancer dataset. We then used the Nimbus scores as inputs to unsupervised clustering using a self-organizing map^12^. We found that this approach enabled us to accurately identify the cell subtypes in the images, which we grouped broadly into cancer, immune, and stromal populations (**Fig. 5d**). Nimbus scores reflected the expected marker staining patterns, with high expression of key lineage defining markers in the appropriate populations such as CD3 in T cells and Ecadherin in Cancer cells (**Fig. 5d**). In addition to these broad lineages, this approach successfully identified more granular subpopulations of cells, such as regulatory T cells (CD3+CD4+FOXP3+) and antigen presenting cells (HLADR+CD11c+).

As a second validation, we used Nimbus to generate per-cell scores for a different MIBI dataset consisting of breast cancer precursor lesions, which we then fed into the same clustering pipeline as above (**Fig. 5e**). Following unsupervised clustering of Nimbus predictions (**Fig. 5e**), we successfully identified the major cell lineages in the image, such as lymphoid cells positive for CD45, and endothelial cells positive for CD31. We also identified granular cell populations such as KRT7+ KRT15+ KRT81+ epithelial cells, CD206+ CD209+ myeloid cells, and KRT14+ SMA+ myoepithelial cells. Across the two datasets, we were able to combine Nimbus with unsupervised clustering to assign 94.66% of the cells to a specific cluster, with only 5.34% of unassigned cells, highlighting the utility of this approach for unsupervised cell population identification.

### Discussion

Robust cell type assignment in spatial data has remained a significant bottleneck in image analysis pipelines. The customization that goes into constructing antibody panels across distinct studies means there is substantial variation in the markers used to define cell types. This prevents the creation of pretrained deep learning models that can generalize beyond the markers they were trained on. Here, we addressed this problem by predicting positivity on a per marker basis, rather than directly predicting cell type. We constructed the Pan-M dataset, containing more than 197 million annotations across 15 million cells. We used Pan-M to train Nimbus, a deep learning model to predict marker positivity one channel at a time. Nimbus can accurately predict marker positivity across the four tissues, three imaging platforms, 10 cell lineages, and 56 markers in Pan-M. These predictions can be leveraged in traditional clustering algorithms to easily identify cell types.

Despite the wealth of spatial information contained within imaging data, nearly all previously developed algorithms to cluster cells in image data operate on the extracted counts per cell, not the actual images. This is in contrast to how experts evaluate the accuracy of clustering, where visual inspection of the underlying images is crucial in order to assign cells to the correct lineage. By training Nimbus directly on a diversity of multiplexed images, we have created an algorithm that much more closely mirrors the workflow of a human expert, but with the scalability inherent to a fully automated deep learning solution.

Although pretrained deep learning models are now available for a wide range of biological image analysis tasks, prior to this work there were none for cell classification. This was not because of an inherent technical barrier, but rather because of how the problem had been posed. Training a model to directly predict cell types based off combinations of markers means that the model must learn which markers are associated with cell types; as a result, study-to-study variation in which markers are used to identify specific cell populations, and which cell populations are being profiled, would necessitate the development of new models. For example, CellSighter^27^ is a recently published deep learning algorithm for cell type prediction, and is one of the only other approaches for cell classification in image data that operates directly at the image level. However, the model must be retrained for each dataset it is applied to. MAPS^21^, another recently published deep learning algorithm for cell classification, likewise must be retrained on each new dataset. Custom models have the potential to generate classifications that precisely conform to the specifics of a given dataset, but the time and effort to accomplish such a task is significant.

Our insight was that clustering image data is more challenging than clustering other types of single cell data not because the cell types themselves are harder to distinguish, but rather because the integrated expression of counts in each cell is noisier due to the spatial nature of the data. As a result, by reframing the task from predicting cell types to predicting marker positivity, we developed a model which exclusively solves the image-specific challenges for cell assignment. Following marker positivity prediction, the single cell imaging data is no more challenging to work with than any other type of data. This means that following single-marker predictions with Nimbus, the data can be clustered using a non-spatial clustering algorithm, taking advantage of the infrastructure that has been developed for other modalities of single-cell data.

Given that Nimbus was trained on Pan-M, a natural question is whether it learned the same biases and errors present in the underlying data. This is a major concern when training models which directly predict cell type, as any inaccuracies in the training data will be baked into the final model. However, the structure of the prediction task we used for Nimbus helps to alleviates this issue. Because we split each cell up into its constitutive channels during training, and only ever predict a single channel at a time,

Nimbus never sees the cell-level biases that exist in the dataset, making it harder for these biases to be learned during training. For example, if one dataset tended to incorrectly label regulatory T cells (Tregs) as helper T cells (CD4T), a model trained to directly predict cell type would learn that same bias. However, because Nimbus was only trained to predict channel scores, rather than cell types, it doesn’t see that the specific combination of markers that define a Treg (CD3, CD4, FOXP3) have been incorrectly labeled a CD4T (CD3, CD4). Instead, it just sees some examples where the silver standard label for FOXP3 is negative, when in fact the true label is positive. Rather than representing a source of systematic bias in the training dataset, this just contributes to the overall error rate of the silver standard dataset; we utilize a training schema which reduces the impact of incorrect labels^33^ to account for this (see Methods).

Although Pan-M and Nimbus represent a major step forward in our ability to analyze multiplexed imaging data, our study has several important limitations. Foremost of these would be the data types that should be analyzed with Nimbus. Nimbus will not perform well on data types not seen during training, such as H&E, immunohistochemistry, or spatial transcriptomics. Additionally, though we attempted to include a wide representation of different tissue types and markers in Pan-M, we were not able to generate an exhaustive training dataset. Given that Nimbus is only as accurate as the data it was trained on, it is likely that Nimbus will not perform well on tissues or markers with very different staining patterns from those present in Pan-M. Finally, since Nimbus does not directly perform downstream cell clustering, it does not solve the issues inherent to current clustering algorithms for high-dimensional data, such as determining the number of distinct clusters, the challenges with identifying rare subpopulations, and variation from non-deterministic algorithms. We anticipate that future work will be able to leverage the template we established here to address many of these shortcomings, setting the stage for further improvements in the robustness, accuracy, and generalizability of biological image analysis algorithms.

## Methods

### Creating the Pan-M Dataset

Our aim was to create a robust computer vision model for multiplexed image analysis, generalizing to diverse cell types, tissue types, and imaging platforms. This required us to create a comprehensive and heterogeneous dataset that encapsulated the variability observed in multiplexed imaging studies. This heterogeneity spanned multiple axes, including four organ systems, three imaging platforms, 10 cell lineages, and 56 markers. Of the three imaging platforms, Vectra and CODEX are both immunofluorescence-based, whereas MIBI-TOF uses mass spectrometry as a readout. We included images from tissue microarrays, as well as whole tissue sections composed of stitched and tiled images, to ensure that the Pan-M dataset was as representative as possible.

To account for diversity introduced by varying computational processing pipelines, the Pan-M dataset incorporates variations in cell segmentation algorithms and cell phenotyping pipelines. Different versions of Mesmer^17^ were used for the MIBI-TOF datasets, whereas the CODEX colon dataset was segmented using the CODEX Segmenter^34^. Cell phenotyping was done via manual gating of individual channels for the two Vectra datasets, using FlowSOM^15^ for the MIBI-TOF decidua dataset, using Pixie^12^ for the MIBI-TOF TNBC dataset, and with STELLAR^20^ for the CODEX dataset. For a full description of the parameters for each dataset, see **Supplementary Table 2**.

### Preprocessing

To harmonize marker intensities across all datasets, the individual channels within each dataset were normalized based on the channel-wide 99.99th percentile of their intensity values. Then, images were resized to 1/4^th^ of their original resolution, to balance computational efficiency without compromising prediction quality (**see Extended Data Fig. 1f**). Images were then cropped to 512^2^ sized tiles with 16 pixels overlap and stored as .tfrecord files for fast loading and model training.

### Automatic creation of silver standard labels

For the MIBI-TOF and CODEX datasets, we generated silver standard labels for individual cells using a semi-automatic approach. For each dataset, we first constructed an assignment matrix that mapped cell types to their specific marker expression patterns. For each cell type, we identify which markers should be positive in that cell, which markers should be negative, and which markers are undefined for that cell type (**Extended Data Fig. 1a-c**). For example, cytotoxic T-cells are mapped to positive marker expression for their lineage defining markers CD45 (lymphocytes), CD3 (T-cells) and CD8 (cytotoxic T-cells), negative marker expression for lineage defining markers of other cell types (e.g. CD14 which is lineage defining for monocytes) and undetermined marker expression for markers whose expression is not uniform across that cell type (e.g. Ki67 as a marker for proliferating cells; see **Supplementary Table 2** for a list of markers that were undefined for all cell types and excluded).

We then validated that the resulting marker profiles for each cell type aligned with the per-cell intensities from the original clustering, and consulted with the original study authors as needed. We used the assignment matrices to map cell types to marker positivity for each marker, and then used the location information of each cell to generate an image-level semantic segmentation mask where the pixels belonging to each cell that was positive for a given marker were positive, and the pixels belonging to cells negative for a given marker were negative.

The two Vectra datasets came with manually assigned per-channel integrated expression thresholds. We used these thresholds to assign cells into marker positive and negative classes for both datasets and generated silver standard semantic segmentations maps.

Finally, we visually examined the silver standard labels of all datasets by comparing them with their corresponding marker images, and identified channels with a high disparity between silver standard labels and marker expression. Since cell type annotations were done with manual gating or unsupervised clustering, we expect that some cells are false positive or negative, thus adding label noise to the dataset. We gauged the amount of label noise by comparing the silver standard annotations against the manually proofread gold standard annotations and report quality metrics in **Fig. 2d**. See **Supplementary Table 2** for a list of markers that were excluded due to poor visual agreement.

### Manual annotation to construct gold standard labels

For three randomly selected images from each of the five studies, we generated gold standard annotations via manual proofreading of the silver standard labels. The silver standard labels were exported to QuPath^35^, and expert annotators looked at each channel and its silver standard annotations individually. The silver standard annotations were systematically corrected by flipping labels from one of (positive, negative, undetermined) to one of (positive, negative, likely positive, likely negative). A consensus mechanism was adopted, where annotators met for weekly discussions to resolve borderline cases and ensure consistent scoring. Following the first round of manual scoring, an independent annotator proofread all annotations a second time to ensure consistency among annotations. We used these gold standard labels only for assessing the accuracy of the model, not for training.

### Nimbus model design

Our goal was to have a model which could implement a coordinate transform from image space (which is confounded by signal intensity, subcellular expression patterns, noise, and other artifacts) to marker confidence scores (which would ideally be free of those confounders and accurately represent the expression of a marker in each cell). Rather than constraining the model to a fixed number or sequence of markers, which would limit general applicability and require retraining, we opted for a design that would compute a score for each marker separately. This design decision allowed for adaptability to different marker sets and enhanced the model’s general applicability across diverse experimental scenarios.

Based on our model design considerations, we opted for a U-Net architecture^31,32^, which takes the normalized tiles of antibody-stained images along with foreground / background cell segmentation maps as the inputs, and outputs pixelwise confidence scores, calculated as the sigmoid of the last layer of the network. These confidence scores capture the chance that a cell is positive for the given antibody in the input image. We compute the average of the per-pixel confidence scores across all pixels in each cell. The U-Net is a convolutional neural network commonly used for biomedical image analysis, due to its ability to capture features at multiple scales. We tested several pre-trained backbones, such as variants of NASnet^36^, EfficientNet^37^ and ResNet^38^, and found that the regular Residual U-Net^32^ achieved the highest accuracy for our task (**Extended Data Fig. 1g**). We also tested whether having additional inputs, such as a nuclei or membrane channel, would increase performance, but saw no difference in accuracy (**Extended Data Fig. 1e**).

### Noise-robust training procedure

Given that the silver standard labels contain errors from the original clustering, we adapted a noise-robust training procedure^33^ originally developed for image classification to help the model avoid overfitting to the erroneous labels in the dataset. An initial model was first pre-trained with a cross-entropy loss on the silver standard labelled dataset using high weight decay to prevent overfitting, but low enough to still enable the model to learn from the data and make predictions. The model was then finetuned by excluding cells from the loss calculation where the model has low confidence or high loss (i.e. confidently disagrees with the noisy labels). Cells were excluded if their loss was above the 85th-percentile of the exponential moving average (EMA) of the loss. This percentile-based EMA threshold is calculated separately for each dataset and marker combination, to ensure that similar proportions of labels for each were retained. In addition to excluding cells with high loss, we specifically selected cells to include using a matched-high confidence selection mechanism^33^. Here, cells were included if the model’s cell-wise predictions and silver standard labels agreed, and the predicted confidence was above 0.9 for positive cells and below 0.1 for negative cells. Using this noise robust training procedure resulted in a modest increase in model accuracy (**Extended Data Fig. 1d**).

### Training details

The fields of view (FOVs) in the datasets were initially split into subsets, with 80% of FOVs assigned to the training dataset and 10% assigned to the validation and test set each. The FOVs that were annotated with the gold standard labels were assigned to the test set. We used the Adam optimizer^39^, a cosine decay learning rate scheduler starting from a learning rate of 3e-4, a weight decay with weight 1e-3, and optimized the model with batchsize 16. To increase the robustness in training, we augmented the data using elastic deformations, flips, rotations, random brightness and contrast, additive gaussian noise and gaussian blurring, implemented via the imgaug library^40^.

The model was first trained with the noise-naïve training procedure for 300,000 steps, then the noise-robust finetuning was applied for 100,000 steps. No early stopping was employed, and the training was continued throughout. The checkpoint with the highest silver standard validation dataset F1-score was selected. The training was conducted using TensorFlow 2.8 on NVIDIA A40 and H100 GPUs.

### Model inference

For inference, we first calculate channel-wise 99.99% pixel intensity percentiles over the whole dataset for normalization. Then, input images are normalized and resized to 1/4^th^ resolution. Additionally, we transform the instance map into a binary representation with eroded boundaries and average predictions over multiple views generated by flipping and 90°-rotations, a technique called test time augmentation that is known to improve performance. Subsequently, post-processing includes the application of inverse augmentations and averaging over test-time augmented predictions. Furthermore, we employ artifact-free tile and stitch inference^41^ for large Fields of View (FOVs) and integrate the Nimbus score per cell segment, storing the channel-wise results in a tabular format which can then be easily used for downstream analysis.

### Validation and benchmarking

To evaluate and benchmark Nimbus confidence scores, we computed the precision, recall, specificity, and F1 scores between Nimbus predictions and the gold standard annotations, as these metrics are robust to class imbalance. Of note, in multiplexed proteomic imaging datasets, class imbalance arises as a result of antibody panel design, where most cells are negative for most markers. A quick guide on interpreting these metrics is as follows: precision represents the share of true positives among all positive predictions, recall indicates the share of true positives among all ground truth positives, specificity reflects the share of true negatives among all negative predictions, and the F1 score combines precision and recall by taking their geometric mean. We benchmarked Nimbus and the silver standard labels using these metrics against the gold standard annotations to assess model accuracy, as well as establish a baseline for the underlying training data. Reported metrics were averaged over the five individual datasets within Pan-M.

### Confounders analysis

We calculated cell-level metrics that we hypothesized might correlate with model accuracy to understand the factors that influence model performance. We used the mutual information criterion^42^ to capture the relationship between possible confounders and errors in the predictions of Nimbus and the silver standard annotations on the gold standard test set. We defined the possible confounders as follows: Cell size was defined as the number of pixels in the segmentation mask of each cell. Marker heterogeneity was defined as the coefficient of variation, which scales the standard deviation by the mean, of the integrated expression for each channel in each FOV. Marker rarity was defined as the share of marker positive cells for a given marker and FOV. Marker sparsity was defined as the number of marker positive cells within a 120 pixel radius of a given cell. Cell density was defined as the total number of cells within a 120 pixel radius of a given cell.

### Cell clustering

To demonstrate the potential of Nimbus in improving cell population identification, we performed unsupervised clustering of cells according to their Nimbus confidence scores using a self-organizing map (SOM)^15^. The SOM is an artificial neural network that aggregates similar cells to one another, resulting in a fixed number of distinct clusters. Here, we “over cluster” the data by specifying a large number (200+) of distinct groups, and then performed hierarchical clustering with manual adjustment to combine these groups together where they represented the same underlying cell type. We took advantage of a workflow and corresponding GUI we recently developed for this task^12^, which significantly speeds up this process and allows for easy manual inspection and correction of the over-clustered data.

The resulting cluster assignments were then manually inspected to identify potential issues with the clustering using both Python scripts and Mantis Viewer^43^. For example, cells in close physical proximity to one another were inspected to ensure that signal spillover did not influence the results, markers with dim expression were double checked to ensure that they were not dwarfed by brighter markers, etc. Following manual inspection, the combination of the over clustered data was modified as necessary to generate the appropriate per-cell assignments.

## Code and data availability

Light-weight and easy to use inference code for Nimbus is available at github.com/angelolab/Nimbus-Inference. Code for preparing the dataset, model training and evaluation is available at github.com/angelolab/Nimbus, and code for figure generation is available at https://github.com/angelolab/publications/tree/main/2024-Rumberger_Greenwald_etal_Nimbus. Our Pan-M dataset can be downloaded at https://huggingface.co/datasets/JLrumberger/Pan-Multiplex.

## Acknowledgements

This work was supported by the IFI program of the German Academic Exchange Service (DAAD) (J.L.R.); NCI CA264307 (N.F.G.) and the Stanford Graduate Fellowship (N.F.G.); NIAID F31AI165180 (C.C.L.) and the Stanford Graduate Fellowship (C.C.L.); NIH grants 5U54CA20997105 (M.A.), 5DP5OD01982205 (M.A.), 1R01CA24063801A1 (M.A.), 5R01AG06827902 (M.A.), 5UH3CA24663303 (M.A.), 5R01CA22952904 (M.A.), 1U24CA22430901 (M.A.), 5R01AG05791504 (M.A), 5R01AG05628705 (M.A.); the Department of Defense W81XWH2110143 (M.A.), the Wellcome Trust (M.A.) and other funding from the Bill and Melinda Gates Foundation (M.A.), Cancer Research Institute (M.A.), the Parker Center for Cancer Immunotherapy (M.A.), and the Breast Cancer Research Foundation (M.A.).

## Ethics declaration

M.A. is an inventor on patents related to MIBI technology (US20150287578A1, WO2016153819A1 and US20180024111A1). M.A. is a consultant, board member, and shareholder in Ionpath Inc. The remaining authors declare no competing interests.

## Contributions

J.L.R., N.F.G., D.K. and M.A. formulated the project. J.L.R. created the deep learning pipeline and trained the models. J.L.R., N.F.G, A.K., S.R.V., C.S., and C.C.L. wrote the software. J.R., H.P. and I.A. ran the cell phenotyping workflows. J.L.R., N.F.G., P.B and C.W. provided manual annotations for the gold standard dataset. R.V, I.N., M.K., and T.V. helped to generate training data. J.L.R. performed the analyses and generated the figures. J.F. revised the figures. J.L.R., N.F.G., and M.A. wrote the manuscript. X.J.W. and D.V.V. provided guidance. N.F.G. and M.A. supervised the work. All authors reviewed the manuscript and provided feedback.

