## Extended Data Figures for "Automated classification of cellular expression in multiplexed imaging data with Nimbus"

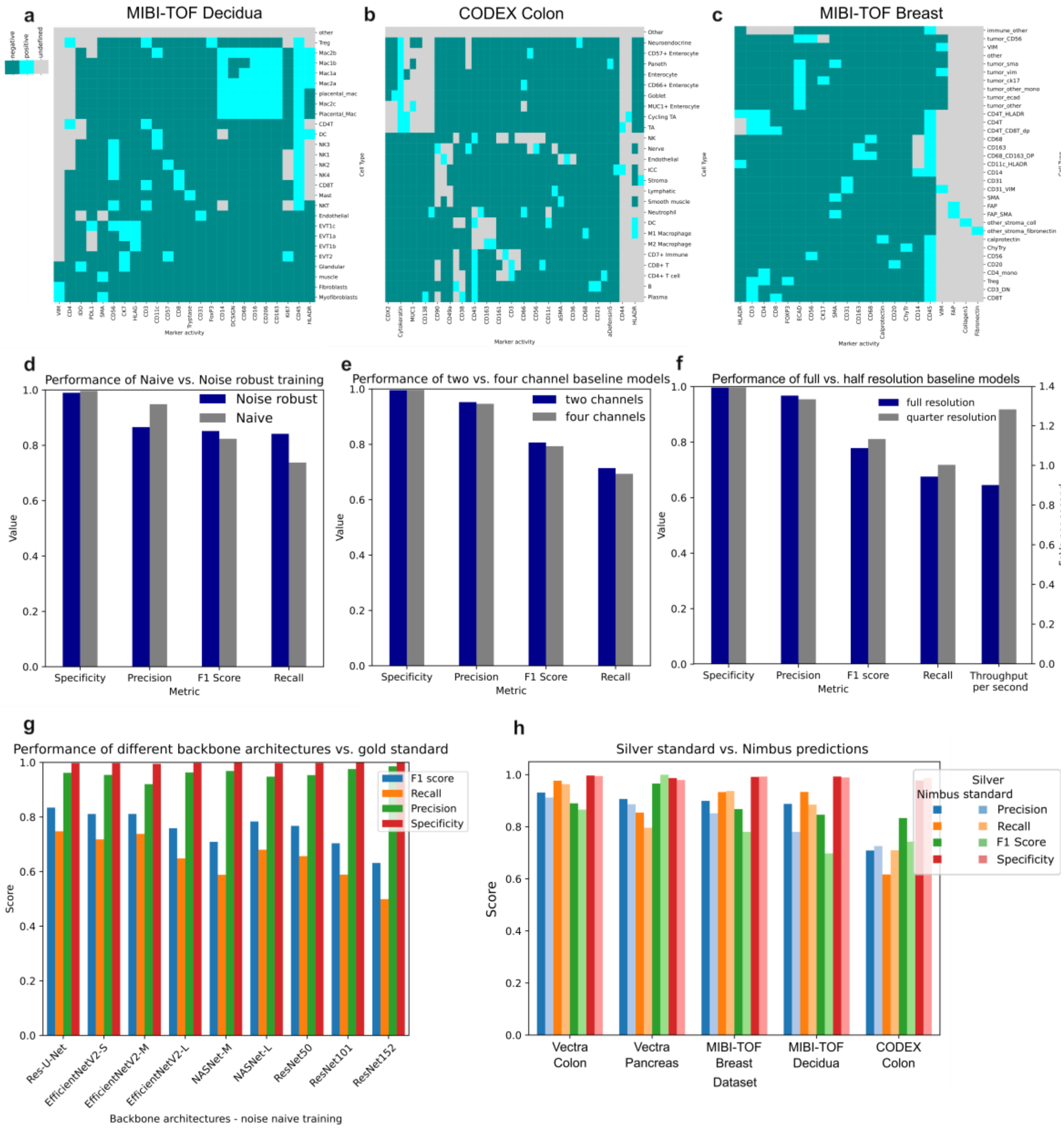


**Extended Data Fig. 1 | Pan-M conversion matrices and NIMBUS accuracy comparisons. a-c**, Conversion matrices used to transform the cell types in the MIBI-TOF Decidua, CODEX Colon and MIBI-TOF Breast to marker positivity estimates. **d**, Performance metrics of naïve vs. noise-robust training procedures. **e**, Performance metrics for models trained with additional nuclei and cell membrane channels (four channels) as input. **f,** Performance metrics for full resolution (20x) vs. quarter resolution (10x) input data. **g**, Performance metrics for different backbone architectures. **h**, Performance metrics vs. gold-standard annotations of the final model split by the datasets in Pan-M.

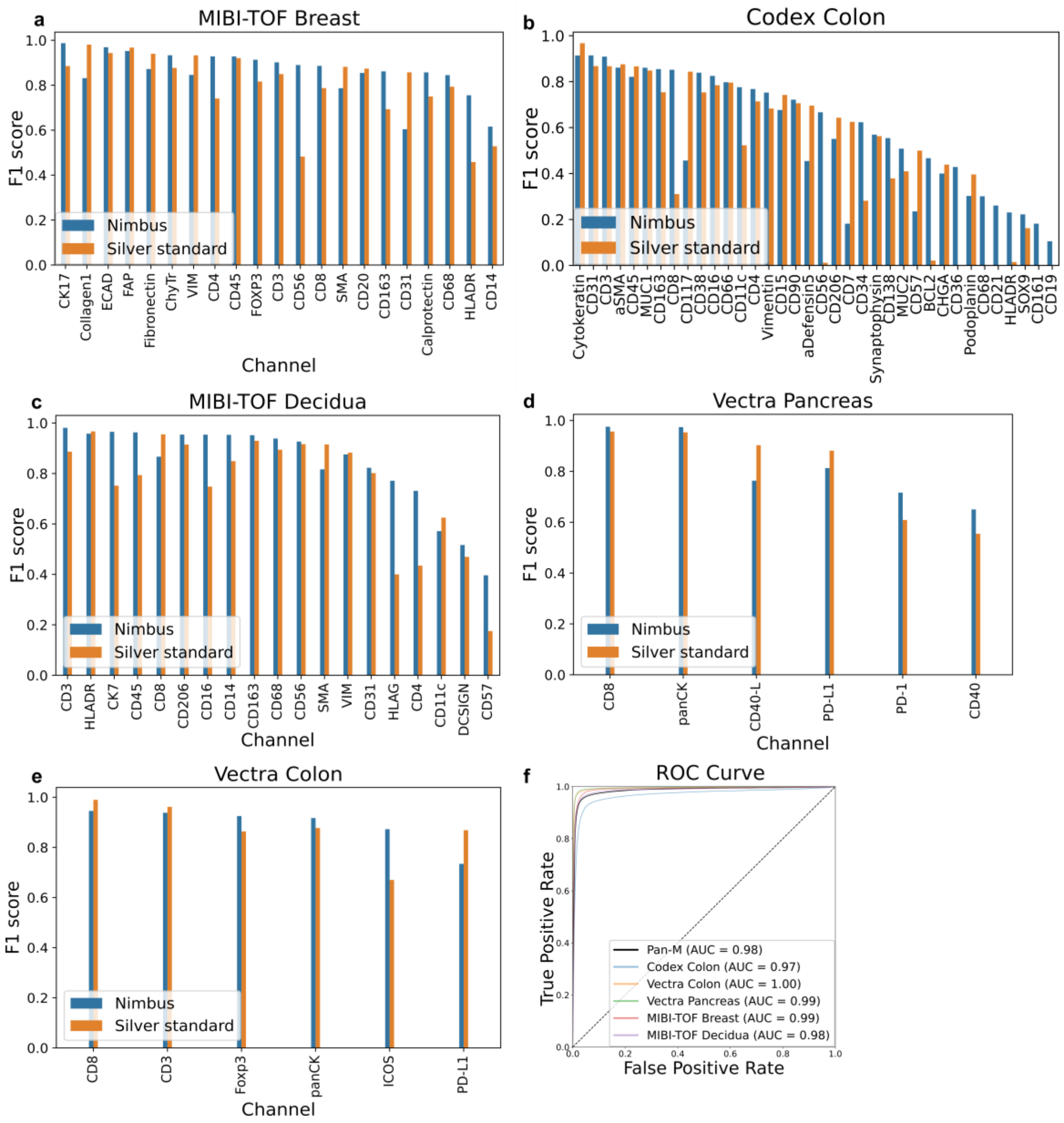


**Extended Data Fig. 2 | NIMBUS F1 score comparison for individual channels. a-e**, F1 scores for the individual channels of the data subsets of Pan-M. **f**, Receiver Operating Characteristic (ROC) curves for the individual datasets and combined Pan-M dataset.
